# An optimized retroviral toolbox for overexpression and genetic perturbation of primary lymphocytes

**DOI:** 10.1101/2021.01.08.425881

**Authors:** Lieve EH van der Donk, Jet van der Spek, Tom van Duivenvoorde, Marieke S ten Brink, Teunis BH Geijtenbeek, Coenraad P Kuijl, Jeroen WJ van Heijst, Louis S Ates

## Abstract

Genetic manipulation of primary lymphocytes is crucial for both clinical purposes and fundamental research. Despite their broad use, we encountered a paucity of data on systematic comparison and optimization of retroviral vectors, the workhorses of genetic modification of primary lymphocytes. Here, we report the construction and validation of a versatile range of retroviral expression vectors. These vectors can be used for the knockdown or overexpression of genes of interest in primary human and murine lymphocytes, in combination with a wide choice of selection and reporter strategies. By streamlining the vector backbone and insert design, these publicly available vectors allow easy interchangeability of the independent building blocks, such as different promoters, fluorescent proteins, surface markers and antibiotic resistance cassettes. We validated these vectors and tested the optimal promoters for *in vitro* and *in vivo* overexpression and knockdown of the murine T cell antigen receptor. By publicly sharing these vectors and the data on their optimization, we aim to facilitate genetic modification of primary lymphocytes for researchers entering this field.

## Introduction

In a scientific era where high-throughput technologies increasingly dictate immunological research, having tools to characterize individual protein functions is still of great importance. The most fundamental molecular biology tools to achieve this are based on genetic perturbation or overexpression of genes of interest. These tools are often developed and optimized over many years in specialized labs. However, for researchers entering this field it can be daunting to select and obtain the most appropriate vectors for their model. These already difficult decisions are hampered by the paucity of published data on systematic comparisons between components of expression systems. We invested significant effort in developing a versatile vector system and performing quality control and optimization experiments. By sharing these data and systems we aspire to facilitate this process for others.

Genetic perturbation of gene expression by deletion or knockdown in eukaryotic cells has been revolutionized in recent decades by the development of RNAinterference approaches and CRISPR-Cas9-based methods. Similarly, the development of high-resolution fluorescent microscopes and novel fluorescent proteins have revolutionized our knowledge of protein localization and trafficking. However, a limiting factor in the genetic manipulation of primary eukaryotic cells is the efficiency of the transfection or transduction method and the stability of the achieved expression. Especially in primary murine and human T cells, it can be challenging to transduce and express large lentiviral constructs, making CRISPR-Cas9 modification of primary T cells technically challenging beyond specialized labs (Hultquist et al., 2016; Schumann et al., 2015; van der Donk et al., 2020). Therefore, genetic perturbation of murine and human T cells is often most readily achieved by expression of optimized microRNAs from gamma-retroviral vectors (Dow et al., 2012; Fellmann et al., 2013; van der Donk et al., 2020). Although gamma-retroviral transduction to achieve gene knockdown or overexpression has been widely used and optimized over recent decades (Dow et al., 2012; Fellmann et al., 2013; Kitamura et al., 2003; Kurachi et al., 2017; Morgan & Boyerinas, 2016), we noted a lack in published literature describing a systematic evaluation of which promoters to use for stable *in vitro* and *in vivo* gene silencing of primary lymphocytes. We set out to select the optimal promoter sequences to stably express proteins and microRNAs of interest in primary T cells *in vitro* and *in vivo*. We constructed a publicly available modular set of vectors, which can be used to express any gene of interest and/or microRNA together with a choice of promoters, linkers, and fluorescent, or surface markers. We tested these different components in primary T cells derived from C7 mice, which express a recombinant T cell receptor (TCR) recognizing a MHC-II-peptide fragment called ESAT6_1-20_, which is derived from *Mycobacterium tuberculosis* (Gallegos, Pamer, & Glickman, 2008; Gallegos et al., 2011, 2016). We expect that this set of easy-to-use optimized vectors will make molecular biology approaches to study primary T cells more widely accessible and adaptable.

## Results

### Development of a versatile vector set for stable gene expression in lymphocytes

We set out to optimize existing (gamma-)retroviral and lentiviral vector backbones for the stable genetic modification of human and murine lymphocytes. We recently reported the systematic comparison of these lentiviral and retroviral backbones for use in human peripheral blood lymphocytes (van der Donk et al., 2020). To achieve those comparisons we streamlined multiple cloning sites to achieve shuttling of identical inserts between the vector backbones. First, we introduced the same multiple cloning site into the retroviral vector backbones pMX and pMY and the lentiviral vector pLenti (Figure 1a) (Kitamura et al., 2003). In parallel, the endogenous SalI site in pMY and the NotI and MluI sites in pLenti were removed. Therefore, all enzyme sites in the MCS are unique cutters in all vectors with the exception of HindIII and SphI in the pLenti backbone. This harmonization of multiple cloning sites allows simple subcloning of any insert from one vector to another, facilitating direct comparisons to select optimal vector systems for specific model systems. To further optimize this vector set, we selected different promoters, which to our knowledge have not been systematically compared in primary lymphocytes (Figure 1b). Finally, we constructed overexpression constructs, which are organized as modules so that individual building blocks can easily be inserted or interchanged (Figure 1b-c). After the promoter, we inserted a first building block, followed by a choice of linkers and a second building block. Selection of an appropriate linker sequence is important and depends on the goal of the experiments. For direct protein fusion of the two building blocks that can for instance be used in protein localization studies, we used a flexible glycine-serine linker (GSGGSG). For production of separate proteins we included either an internal ribosome entry site (IRES) sequence or a P2A sequence. The IRES sequence is longer and may therefore reduce expression levels of the inserts, but it has the advantage that no remnants of the sequence will be translated (Pestova, Hellen, & Shatsky, 1996). In contrast, the short P2A sequence induces efficient ribosome skipping that leads to separate translation of the two building blocks. However, after this "cleavage" the majority of the 2A amino acids remain on the C terminus of the block 1 protein and the terminal proline becomes part of the block 2 protein (Liu et al., 2017). These extra amino acids may interfere with the correct localization or function of certain proteins. Please note that while 2A fusion results in equimolar amounts of the two proteins, IRES fusion may result in a 3:1 ratio of the two proteins (Goedhart et al., 2011). To select for, or monitor the expression of, the gene-of-interest we selected a wide range of reporter genes. These include genes encoding the fluorescent proteins mTurquoise2, eGFP, mVenus and mCherry, antibiotic selection cassettes against puromycin or blasticidin and non-immunogenic murine cell surface markers Ly6G and CD90.2 (Figure 1c, Supplemental figure 1) (Cormack, Valdivia, & Falkow, 1996; Goedhart et al., 2012; Kremers, Goedhart, Van Munster, & Gadella, 2006; Shaner et al., 2004).

**Figure 1:**
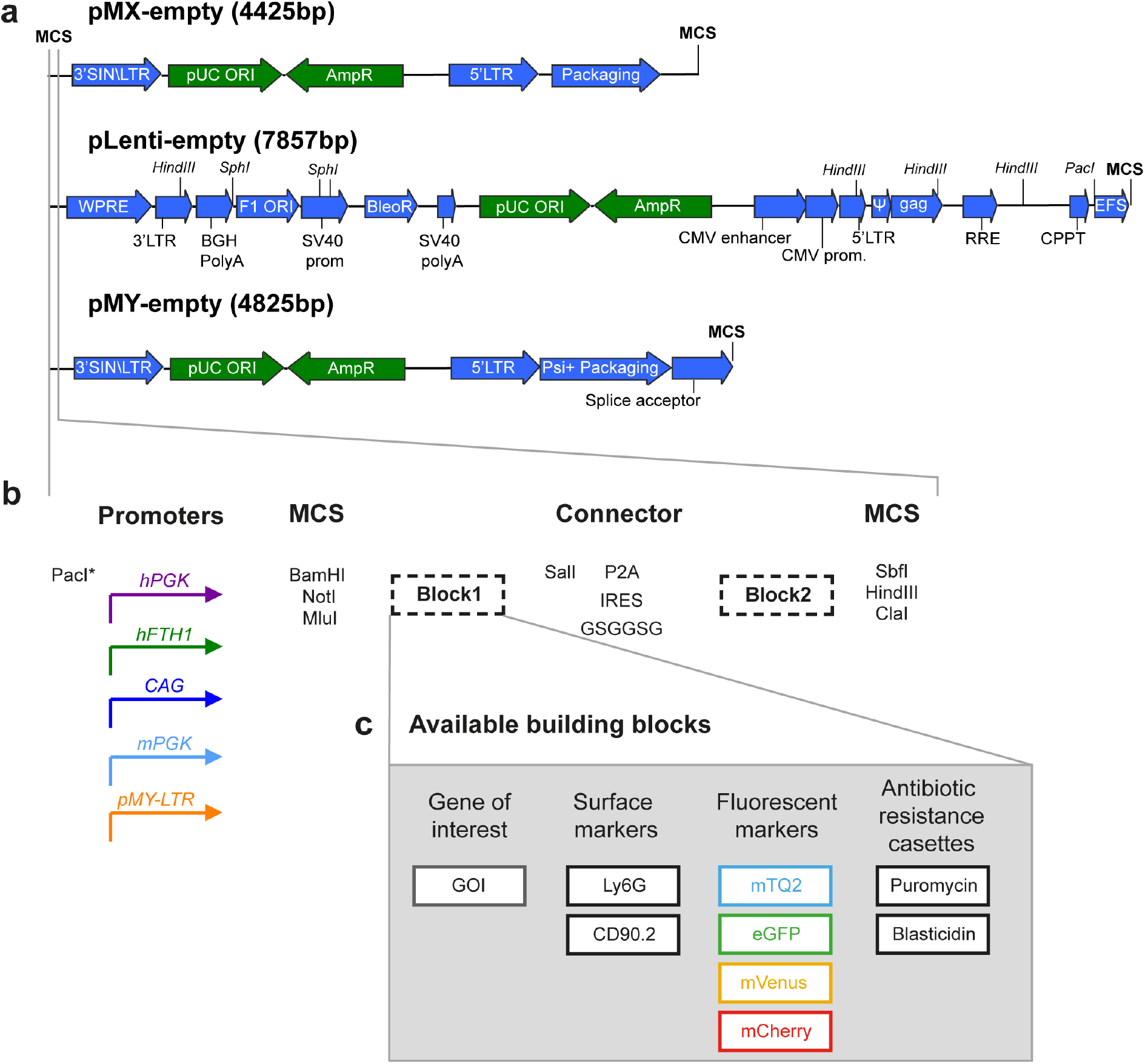
Vector backbones and design. a) Vector backbones of pMX, pMY, and pLenti. Green arrows indicate elements needed for bacterial replication; blue arrows are part of the retroviral genome. b) Organization of the versatile expression cassette used in all three vector backbones. A choice of five different promoter sequences (colored arrows) was assessed for optimal expression *in vitro* and *in vivo*. Note that the lentiviral and pMY backbones already include the EFS/EF1α and LTR promoters respectively and therefore include the PacI site in the MCS after the promoter. Overexpression constructs are designed to express the gene of interest in either block 1 (for C-terminal labelling) or block 2 (for N-terminal labelling). These building blocks are separated by an in frame SalI restriction site and a choice of IRES, P2A peptide, or GSGGSG linker to create a fusion protein or equimolar separately produced proteins. c) Available building blocks include surface markers of murine Ly6G, CD90.1 (Thy1.1) and CD90.2 (Thy1.2), fluorescent proteins mTurquoise2, eGFP, mVenus and mCherry and the antibiotic resistance cassettes conferring resistance to either puromycin or blasticidin.

### Promoter selection by overexpression of a large construct in primary murine T cells *in vitro*

Gene expression from retroviral vectors is generally more stable when the insert is limited in size (Addgene, 2017). To get a stringent read out of which promoters in our expression system are most stable and efficient in murine T cells, we selected a construct which in our hands was relatively challenging to express at high levels. This construct, consisting of the four intracellular components of the T cell receptor (TCR)/CD3 complex consists of codon optimized murine genes encoding CD3γ, CD3δ, CD3ε, TCRζ (CD247) and the marker CD90.2. These genes were separated by the self-dissociating peptides T2A, F2A, E2A and P2A respectively (Liu et al., 2017). This insert was expressed from pMX vectors with the promoters hPGK, hFTH1, CAG, or mPGK and from the pMY vector with its native LTR promoter (Addgene vectors #163334-8). We transduced CD4^+^ T cells isolated from C7 mice, which express a recombinant TCR and therefore all recognize the same epitope derived from *M. tuberculosis* (Gallegos et al., 2008). This transduction resulted in high initial transduction efficiencies, ranging from 60% for the constructs under control of the CAG promoter to almost 100% for the pMY based vector (Figure 2a). However, in line with our previous experience regarding the unstable expression of such a large construct, the percentage of CD90.2^+^ CD4^+^ T cells diminished markedly over time for all constructs. Where these levels approached 0% for vectors under control of the CAG and hFTH1 promoters, the pMY-LTR and hPGK based vectors seemed most stable (Figure 2a). These trends were confirmed by investigating the expression level of CD90.2 within the CD90.2^+^ cells (Figure 2b). Quantification of TCR/CD3 subunit overexpression was hampered by expression of the endogenous TCR on these CD4^+^ T cells. However, a clear overexpression of the different TCR subunits TCRζ, TCRβ and CD3ε could be detected for all vectors 2 days post transduction and was most pronounced with the pMY vector (figures 2c-e). Overexpression of individual TCR/CD3 proteins could only be detected when the CD3 construct was expressed in pMY at day 5 post transduction and was undetectable in all conditions beyond that timepoint (figures 2c-e). Our data therefore suggest that the pMY vector with its native LTR promoter is the most efficient and stable way to overexpress large protein constructs in murine CD4^+^ T cells *in vitro*.

**Figure 2:**
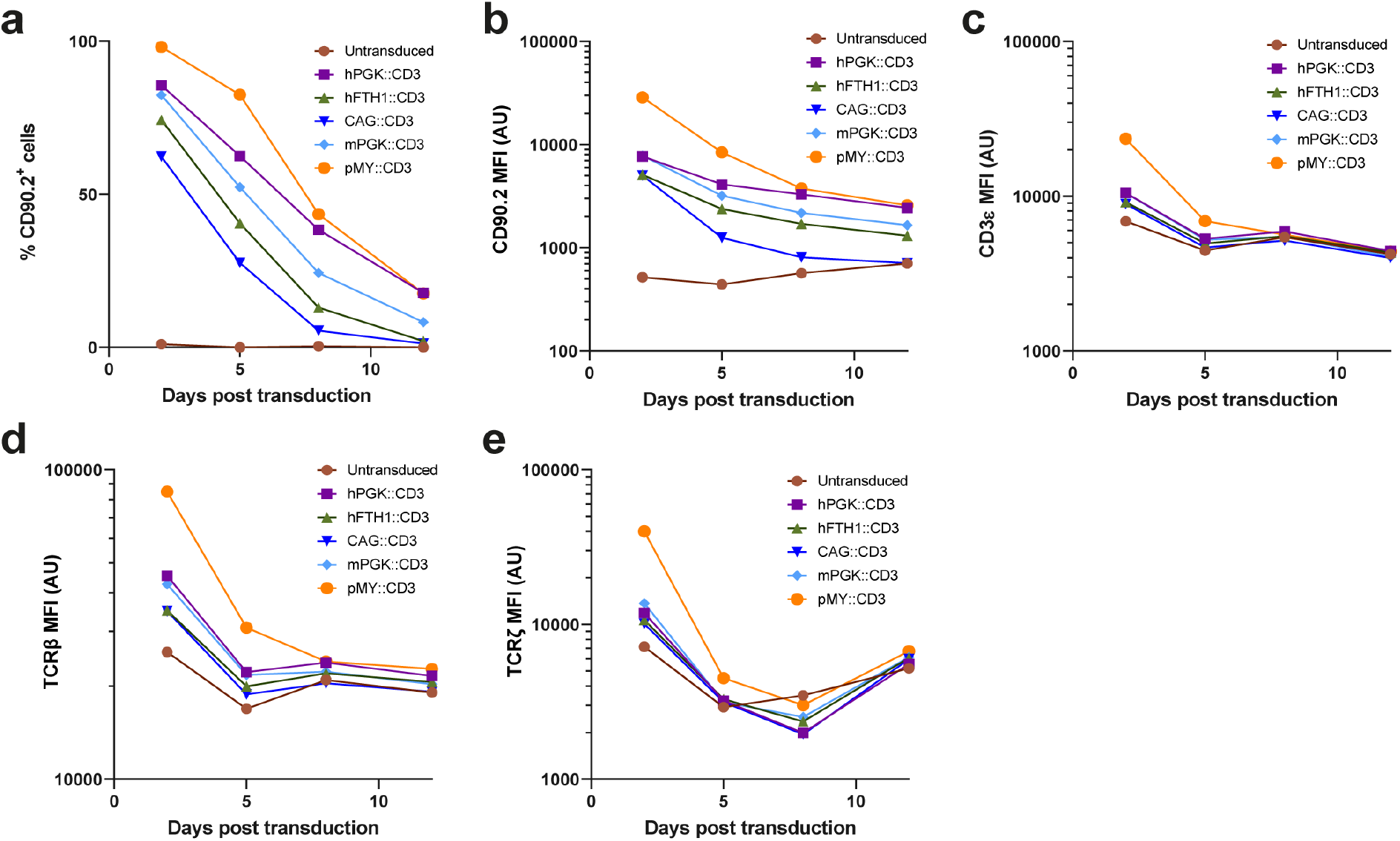
Testing promoter fidelity *in vitro* by overexpression of an oversized construct. Murine C7 splenocytes were transduced with the indicated constructs 2 days after activation with ESAT-6_1-20_ peptide in the presence of IL-12. Expression of CD90.2 (a, b), CD3ε (c), TCRβ (d) and TCRζ (e) was measured 2, 5, 8 and 12 days after transduction by flow cytometry within the viable CD45^+^, CD90.1^+^, CD4^+^ cells. Cells were maintained in the presence of IL-2 after transduction, which was replaced with IL-7 at 8-days after transduction.

### Vectors for protein localization in T cells by multicolor confocal microscopy

Having established pMY as the most efficient vector to overexpress proteins in murine T cells *in vitro*, we constructed a range of these vectors where any gene of interested can be cloned in frame to a C-terminal GSGGSG linker followed by a choice of fluorescent proteins with different excitation and emission spectra (Figure 1b-c, Supplemental figure 1). For proof of concept, we selected murine *lyz2*, which is of interest to our ongoing research. *Lyz2* was cloned N-terminally of genes encoding either mTurquoise2, mVenus or mCherry (Addgene vectors #163347-9) (Goedhart et al., 2012; Kremers et al., 2006; Shaner et al., 2004). It should be noted that mVenus and GFP have partly overlapping fluorescent emission and excitation spectra and it is therefore not advised to use these in combination. We selected mVenus over eGFP for its excellent brightness and monomeric nature (Lambert, 2019). We transduced C7 CD4^+^ T cells with these single vectors, or combinations thereof. Cells were fixed and mounted 2 days after transduction and were analyzed by confocal microscopy (Figures 3a-b). The lyz2-fluorophore combinations localized in cellular compartments that resemble either lysosomes or secretory granules, independently of which fluorophore was used (Figure 3a). Because of the three fluorophores' different spectra, these could be easily imaged without significant background in the other channels (Figure 3a). Double transduction with two viruses simultaneously (Lyz2-mTQ2 + Lyz2-mCherry, or Lyz2-mVenus + Lyz2-mCherry) resulted in a full colocalization of fluorescent compartments, without signal in the remaining channel (Figure 3b). These data suggest that these vectors can be readily used for protein colocalization studies. Therefore, we set out to provide further proof of concept with simultaneous transduction of three different retroviral vectors, Lyz2-mTQ2, GFP-Rab27a and Mpeg1-mCherry (Figure 3c). Different localization of the three fluorophores was observed in the imaged cells, indicating that indeed, triple transduction with our multicolored pMY vectors is a viable approach to study intracellular protein localization in primary murine CD4^+^ T cells.

**Figure 3:**
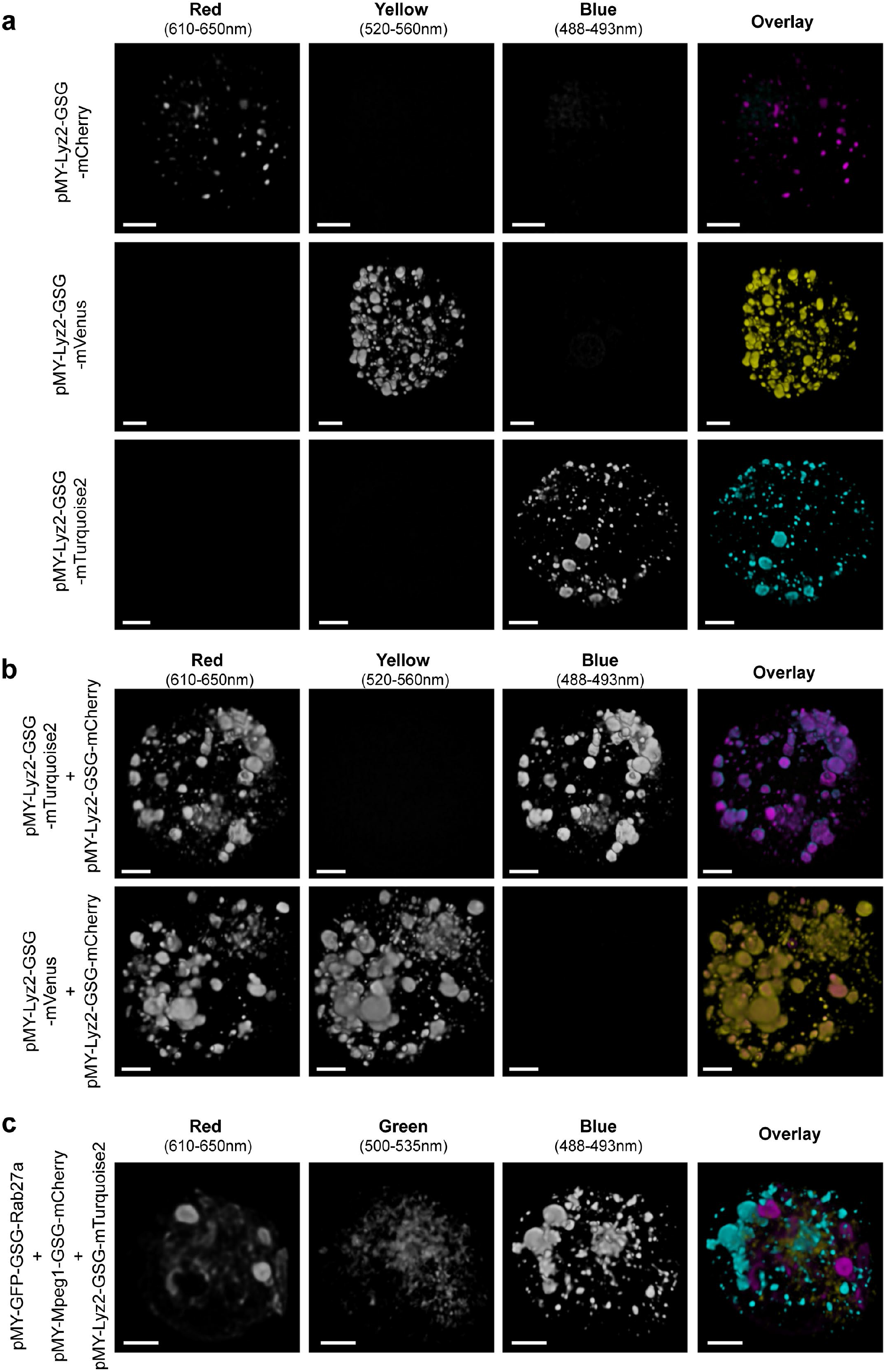
Validation of vectors encoding fluorescent proteins for cellular localization studies. Murine C7 CD4^+^ T cells were transduced with the indicated vectors 2 days after activation. Cells were fixed and mounted to be imaged by confocal microscopy 2 days after transduction. a-c) 3D rendering of Z-stacks with single channels depicted in greyscale and overlays of channels in artificial Cyan, Yellow and Magenta. a) Transduction with single vectors expressing Lyz2, labeled by either mTurquoise2, mVenus, or mCherry shows similar subcellular localization independent of the fluorophore used and minimal spectral overlap in the other channels. B) Co-transduction of two viral vectors was an effective approach to obtain cells expressing both constructs and led to full colocalization of the differently labeled proteins. c) An example of triply transduced C7 CD4^+^ T cells co-expressing Lyz2-mTurquoise, GFP-Rab27a and Mpeg1-mCherry. Scale bars represent 2 nm.

### Assessment of optimal promoters for retroviral mediated knockdown in primary murine T cells *in vitro*

Besides creating vectors for overexpression studies, we were also interested in creating a range of retroviral and lentiviral vectors that can be used to knockdown genes of interest by expressing microRNAs (Dow 2012). To select the optimal constructs for *in vitro* and *in vivo* microRNA-mediated knockdown, we tested the surface marker expression levels and knockdown efficiency of a range of TCRζ microRNA constructs. Since TCRζ is the limiting component for TCR surface expression, knockdown of TCRζ was expected to result in full loss of TCR/CD3 surface expression. First, we selected five target sequences for mouse *Cd247*/TCRζ from the genetic perturbation platform (https://portals.broadinstitute.org/gpp/public/) (Supplemental table 1), cloned corresponding antagomirs into a pMX-based vector, and tested for highest knockdown efficiency. The first target sequence (TRCN0000068158) was found to be most efficient in knockdown of TCRζ and ensuing reduction of surface TCRβ levels (Data not shown). The antagomir targeting this sequence was cloned in pMY, or in pMX vectors with four different promoters (hPGK, hFTH1, CAG and mPGK), followed by the surface marker CD90.2 (Figure 4a, Addgene vectors: 163324-8). As a non-target microRNA control, we selected a specific antagomir targeting the Rluc gene encoding *Renilla* luciferase (Supplemental table 1, Addgene vectors 163329-33).

**Figure 4:**
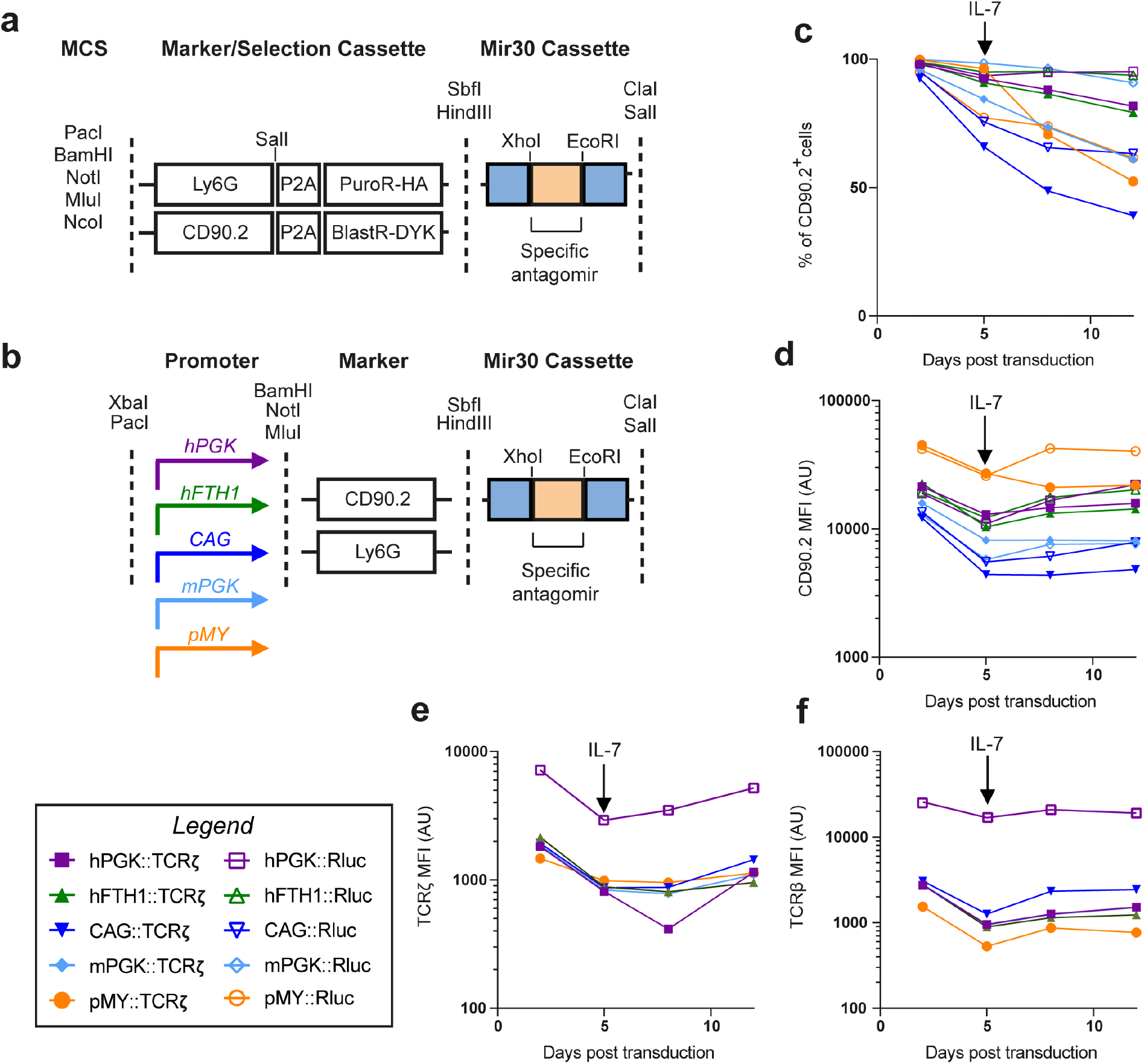
Design and *in vitro* validation of microRNA knockdown vectors. a) Design of vector inserts containing resistance cassettes to puromycin (PuroR), or blasticidin (BlastR). The vectors contain surface markers Ly6G or CD90.2, as well as an HA/FLAG epitope tag on the antibiotic resistance genes, to assess the transduction efficiency by flow cytometry. b) Design of microRNA knockdown vectors suitable for *in vivo* experiments. The microRNA cassette is placed after the surface markers CD90.2 or Ly6G. Four different promoters were assessed in the pMX backbone and compared to the pMY vector with its LTR promoter. c-f) Murine C7 CD4^+^ T cells are activated and cultured *in vitro* and transduced with the indicated vectors containing a microRNA targeting *cd247* (TCRζ, filled symbols) or the *renilla* firefly luciferase (Rluc, empty symbols) gene as a non-target control. Transduction efficiency and stability were assessed by measuring the percentage of CD90.2^+^ cells within the CD45^+^, CD90.1^+^, CD4^+^ cells (c) and the CD90.2 expression levels within CD90.2^+^ cells (d) at 2, 5, 8 and 12 days after transduction. Knockdown efficiency was assessed by measuring intracellular TCRζ (e) and surface TCRβ (f) within the CD90.2^+^ population.

We compared *in vitro* TCR-knockdown efficiency by transducing activated primary murine C7 CD4^+^ T-cells with the respective retroviral knockdown constructs. We cultured C7 CD4^+^ T-cells in the presence of IL-2 until 5 days after transduction, at which point it was replaced with IL-7 to allow long term *in vitro* culture of the T cells. The proportion of C7 cells expressing the retroviral construct was measured by CD90.2 staining and flow cytometry. The highest and most-stable expression of the knockdown construct was achieved by the constructs containing the hPGK and hFTH1 promoters (Figure 4c). However, the expression level of CD90.2 within the CD90.2^+^ cells was highest in the cells transduced with the pMY vectors (Figure 4d). Expression from the CAG or mPGK promoters seemed to be the least efficient. Although differences in CD90.2 expression levels were pronounced between constructs, the knockdown efficiency of TCRζ (Figure 4e) and surface TCRβ (Figure 4f) were similar between the different constructs. This likely indicates that a plateau is reached for this efficient antagomir. Both the proportion of CD90.2^+^ cells (Figure 4c) and the CD90.2 expression within those cells (Figure 4d) in the pMY-transduced cells seemed to fall considerably after the addition of IL-7 to the culture medium. Therefore, we repeated the experiment with a culture maintained on IL-2, which was discontinued at 8 days post transduction. Similar to the first experiment, the percentage of cells that expressed the construct was highest under control of the hPGK and hFTH1 promoters (Supplemental Figure 2a). However, these were now closely followed by pMY, which did not exhibit the marked decline of CD90.2^+^ cells, previously observed after addition of IL-7. Expression levels of CD90.2 and reduction of surface TCRβ were highest in the constructs under control of the pMY-LTR and hPGK promoters and lowest under control of the CAG promoter (Supplemental Figure 2a-d). Together we conclude that for *in vitro* gene-silencing the pMY-LTR, hPGK and hFTH1 promoters are most efficient. Of these, pMY seemed to facilitate slightly higher expression levels leading to the most considerable reduction in surface TCRβ, whereas expression was more stable from hPGK and hFTH1 in the presence of IL-7. Because expression from pMY achieved the highest expression levels, but the expression of this vector was less stable, we also created pMY based knockdown vectors with puromycin and blasticidin resistance cassettes as well as the CD90.2 and Ly6G surface markers (Figure 4a, Addgene vectors 163340-1). These are our vectors of choice when knocking down genes *in vitro* in primary murine or human lymphocytes and can be used for single knockdown as well as for simultaneous knockdown of two genes of interest as we recently demonstrated in van der Donk et al. (van der Donk et al., 2020).

### Comparison of promoters for retroviral mediated knockdown in CD4+ T cells *in vivo*

In parallel with the *in vitro* assessment of expression levels and stability described above, we also assessed these characteristics *in vivo*. The same transduced cells depicted in figure 3c-f were administered to wild-type C57Bl/6 mice via intravenous injection of 2*10^6^ transduced CD4^+^ T cells 2 days after transduction (*i.e*. 4 days after T cell activation). The number and characteristics of the transferred C7 CD4^+^ T cells was followed over time by tail vain bleeds at the indicated intervals. The percentage of CD90.1^+^ cells (C7 T cells) of total leukocytes (defined here as CD45^+^ cells) was comparable for all promoter constructs except for the pMY vector which was markedly lower from day 10 onwards (Figure 5a). The level of CD90.2 expression was consistently lowest for constructs under control of the CAG and mPGK promoters. For the pMY vector, CD90.2 marker gene expression was initially high, but dropped significantly at 22 days post activation (Figure 5b). In contrast, the construct under control of the hPGK promoter was initially expressed at relatively low levels but this expression level was stable and even increased over time (Figure 5b). The efficiency of TCRζ knockdown was assessed by staining intracellular TCRζ as well as surface TCRβ and CD3ε (Figure 5c-e). Although differences between the constructs were modest, knockdown was clearly impaired at the latest timepoint when under control of the pMY promoter. Trends for the most efficient phenotypic knockdown closely resembled expression levels of CD90.2, with the most efficient and most stable knockdown being achieved under control of the hFTH1 promoter (Figure 5c-e). At the termination of the experiment, 22 days after T cell activation, we collected and homogenized lungs and spleens of recipient mice to assess knockdown efficiency in resident CD4^+^ T cells within these target tissues. Spleen and lung phenotypes closely resembled each other and showed considerably lower percentages of CD90.2^+^ cells when expressed from the pMY vector (Figures 5f, 5i). CD90.2 expression levels within the CD90.2^+^ cells were highest for constructs under control of hFTH1 and hPGK promoters although the latter was quite variable between individual animals (Figures 5g, 5j). Surface levels of TCR components did not markedly differ between the constructs expressed from the different pMX vectors, but decreased to control levels (Luciferase antagomir) with the pMY based vector (Figure 5h, 5k). Taken together, we conclude that the hFTH1 promoter is likely the prime candidate to achieve stable and high expression levels *in vivo*. These combined data indicate that our vectors can be used for efficient gene knockdown in primary murine CD4^+^ T cells and that vector expression and knockdown efficiency can be maintained over relatively long timespans allowing for *in vivo* experiments in the context of for instance *M. tuberculosis* infection (Gallegos et al., 2008, 2011, 2016).

**Figure 5:**
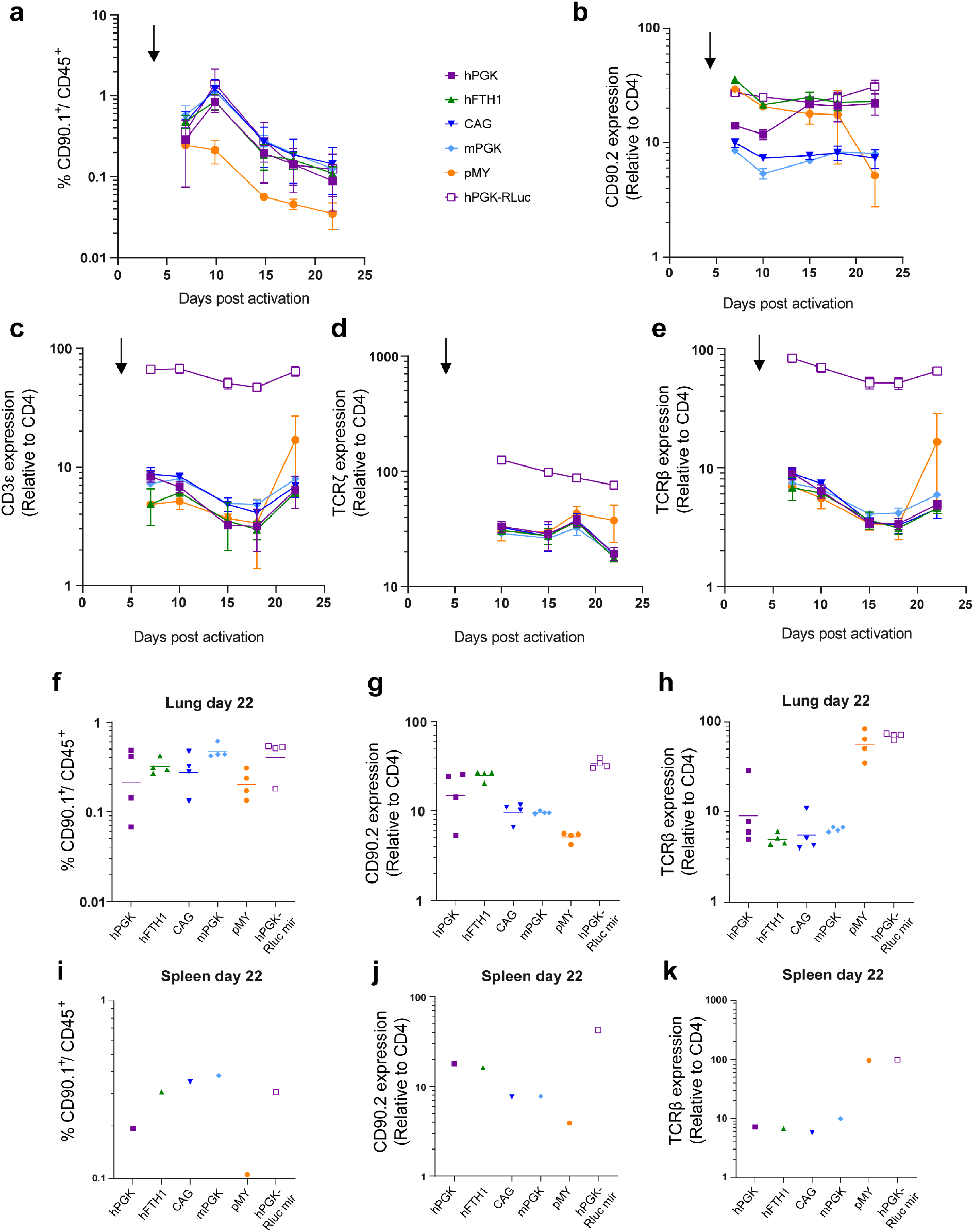
*In vivo* validation of microRNA knockdown vectors. Murine C7 CD4^+^ T cells were activated and transduced before being adoptively transferred to wild-type C57Bl/6 mice by tail-vein injection 3 days post transduction (black arrow). a-e) A maximum of 50 μl of blood was obtained from the mice at the indicated time points by tail-vein bleeds to investigate by flow cytometry. n=4 per group with some exceptions were adoptive transfer was unsuccessful. Individual values are depicted in supplemental figure 3. a) The number of remaining C7 T cells was measured by assessing the percentage of CD90.1^+^ cells within the CD45^+^ cells. b) The level of CD90.2 expression within the CD90.1^+^ cells was normalized to CD4^+^ expression and used as a read-out of expression stability of the vector. c-e) Knockdown efficiency was measured by intracellular staining of TCRζ (d) and surface expression of CD3ε (c) and TCRβ (e). At the end of the experiment, mice were killed and lungs (f-h) and spleens (i-k) were homogenized to assess the percentage of remaining C7 cells (f, i), vector expression stability within the C7 cells (g, j), and knockdown efficiency of the TCR (h, k).

## Discussion

Laboratories that specialize in genetic modification of eukaryotic cells accrue and optimize their tools over the years. Although novel tools that push technical boundaries are often well described, tools that are the "workhorses" of genetic modification and especially data on their optimization or limitations, can be hard to find in published literature. We have developed a versatile toolbox of retroviral vectors, which can be readily used to induce or impair the expression of genes of interest in the context of a wide range of markers. We optimized our vectors allowing efficient cloning of building blocks in different vector backbones and we have tested the optimal promoter sequence for overexpression and genetic perturbation in primary murine lymphocytes *in vitro* and *in vivo*. These same vectors have been recently compared and used in primary human lymphocytes, further confirming their versatility (van der Donk et al., 2020).

It should be emphasized that these vector backbones, as well as the different promoter sequences, surface and fluorescent markers and shRNA-miR building blocks are not in themselves novel and are the result of decades of research by others (Chang, Marran, Valentine, & Hannon, 2013a; Chertkova et al., 2017; Cormack et al., 1996; Dow et al., 2012; Fellmann et al., 2013; Goedhart et al., 2012; Kitamura et al., 2003; Kremers et al., 2006; Kurachi et al., 2017; Lambert, 2019; Naviaux, Costanzi, Haas, & Verma, 1996; Shaner et al., 2004). Furthermore, some researchers may prefer other methods to genetic perturbation based on CRISPR-Cas9 approaches (Jinek et al., 2012; Roth et al., 2018). However, these approaches have been particularly challenging to implement for primary murine and human lymphocytes. Although these difficulties can be circumvented by advanced methodology such as electroporation with purified Cas9 protein, implementing this requires costly reagents, or a specialized laboratory with a wide range of expertise (Hultquist et al., 2016; Roth et al., 2018; Schumann et al., 2015). We were similarly unable to transduce primary murine or human T cells efficiently with retroviral and lentiviral Cas9 expression vectors and have therefore opted for the optimized shRNA-miR strategy of genetic perturbation instead (van der Donk et al., 2020). It should be noted that research efforts to optimize microRNA design have markedly improved this technique over the last decade and knockdown efficiencies of >90% were in our experience often achievable with the system employed here (Chang et al., 2013a; Dow et al., 2012; Fellmann et al., 2013).

Since we were unable to find any systematic comparison of the most efficient and stable promoters to express proteins and miRNA's in murine lymphocytes, we decided to perform these comparisons in this work. We find that the pMY vector backbone with its LTR promoter was the strongest promoter for *in vitro* experiments and therefore this is our promoter of choice in short-term *in vitro* experiments. We created versions of the pMY vectors with antibiotic resistance cassettes to further circumvent reduction of expression during *in vitro* culture. The silencing of pMY-based expression is likely due to the presence of IL-7 *in vivo* and in our extended *in vitro* culture conditions (Tsunetsugu-Yokota et al., 2016). Based on our *in vivo* experiments we conclude that the hFTH1 promoter is an excellent candidate to drive stable expression in adoptively transferred murine lymphocytes *in vivo*. The hPGK promoter may be the best choice when a single promoter for high and stable expression is needed for both *in vitro* and *in vivo* experiments (Adra, Boer, & McBurney, 1987). Surprisingly, we consistently observed higher expression driven by the hPGK promoter than the mPGK promoter, which could be due to a lack of endogenous repressors. This may similarly explain the efficiency of the hFTH1 promoter. Although the data on promoter performance provided here may be important for other researchers’ experimental design, other experimental models may require independent optimization.

When these vectors are used for long-term *in vivo* experiments, such as the adoptive T cell transfer experiment described here, extra care should be taken in their choice and design. The overexpression of heterologous proteins, such as antibiotic selection markers and fluorescent proteins can lead to the development of adaptive immune responses against these components (Stripecke et al., 1999). Such immune responses could result in rejection of the adoptively transferred cells and thereby invalidate potential research findings. Therefore, we advise to only use these antibiotic and fluorescent selection markers *in vitro*, or in short-term *in vivo* experiments and opt for the Ly6G and CD90 surface markers for long-term *in vivo* experiments. Similar care should be taken to vector design on a molecular level, especially in the context of overexpression of a gene of interest. Firstly, the gene of interest should be investigated whether N-terminal or C-terminal tagging is expected to interfere with the protein product's correct localization and function. When a C-terminal tag is not expected to have negative consequences, this may be preferable for reporter constructs, since the protein of interest can be expected to be produced in at least equimolar amounts as the reporter. Therefore, we have focused on C-terminal reporter constructs. In cases where it is expected or experimentally found that both N-terminal and C-terminal tagging interfere with protein function the IRES-sequence can be used to create transcriptional fusion of separate protein products.

Together, the vectors presented here form a versatile “starter set” for researchers with the ambition to apply molecular biology approaches to validate their research. We sincerely hope that sharing these vectors and the data regarding their optimization will aid researchers in immunology to apply these molecular techniques to their research.

## Acknowledgements

The authors would like to thank Joachim Goedhart for providing plasmids, insightful discussion and proofreading of the manuscript. We thank the microscopy twitter community for providing feedback shaping figure 3. We thank Chiara Montironi and Eric Eldering for sharing plasmid pMSCV-IRES-mCherry. We thank Tom van der Poll for sharing expertise and resources. We thank the imaging core facility of the AmsterdamUMC for use of confocal microscope and technical assistance.

LEHvdD, JvdS, JWJvH and LSA were supported by NWO-VIDI grant 91717305 to JWJvH). LSA was furthermore supported by a PostDoc stipend of the Amsterdam Infection and Immunity Institute.

## Ethics statements

All animal experiments have been approved by the Dutch central committee for registration of animal experiments (CCD: Project DIX298) as well as the local committee of the AmsterdamUMC (formerly AMC, Work protocol DSK298).

## Supplemental Figures

**Supplemental Figure 1:**
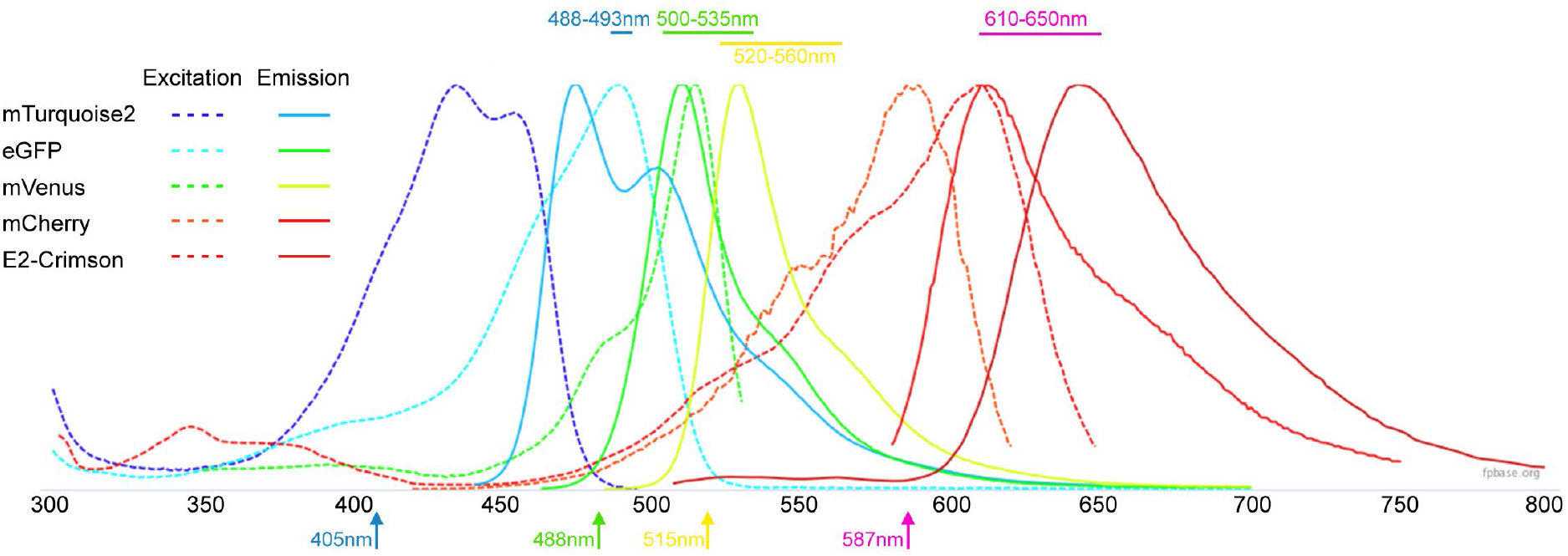
Excitation and emission spectra of the used fluorescent proteins. The X-axis depicts the wavelength in nm; Y-axis depicts relative fluorescent intensity for each fluorophore. Primary investigations with E2-Crimson were not promising and this protein was therefore not included in the vector set. Not that excitation and emission spectra of eGFP and mVenus are quite close and therefore these proteins should only be used together with that knowledge considered. Colored arrows at the bottom indicate used excitation wavelengths. Bars and text at the top indicate acquisition wavelengths used for mTQ2, GFP, mVenus and mCherry respectively. Figure was created with fpbase.org spectra-viewer tool (Lambert, 2019).

**Supplemental Figure 2:**
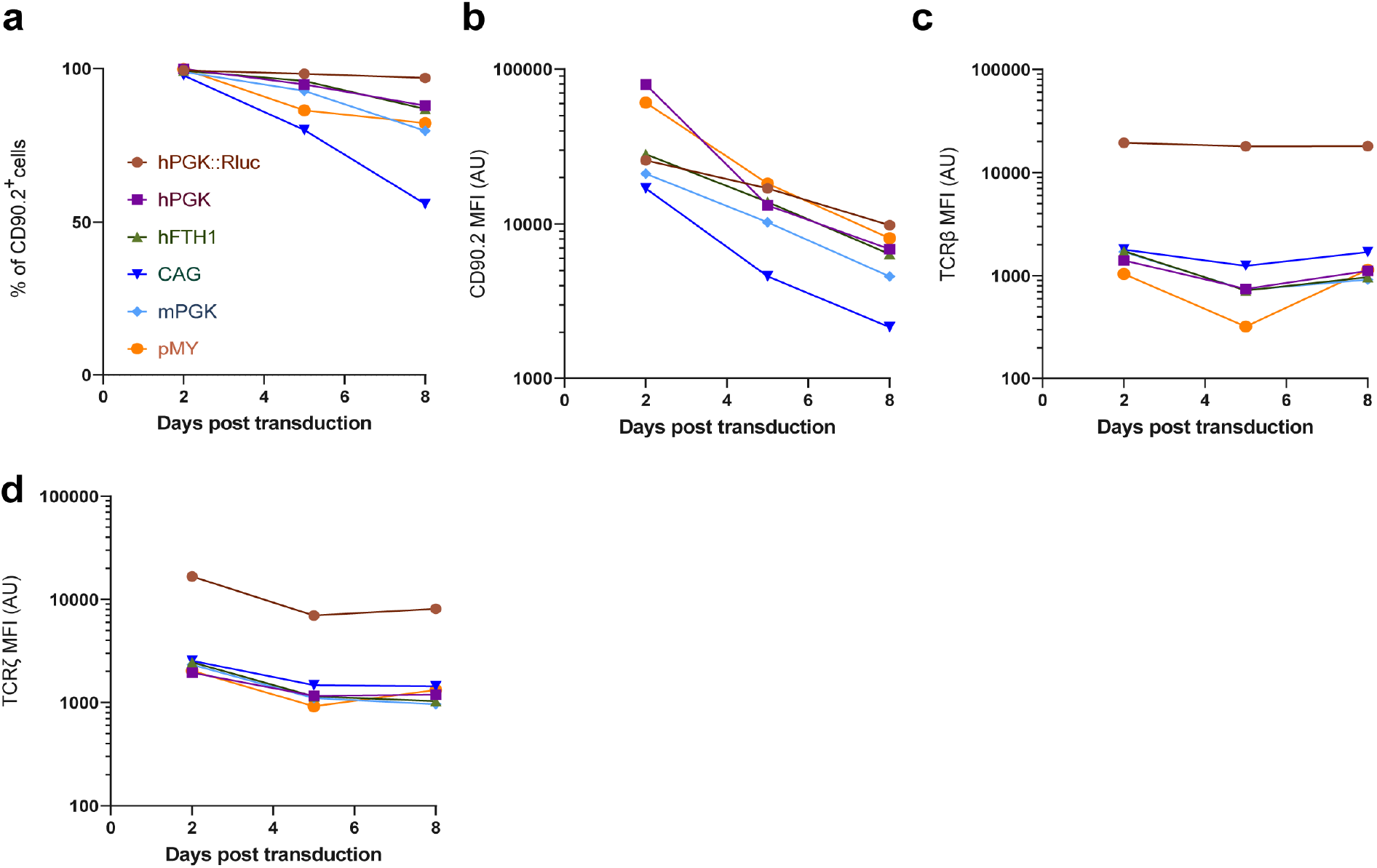
Vector stability and expression *in vitro*, without the addition of IL-7. The experiment in Figure 4c-f was repeated, but cells were maintained constantly on IL-2 instead of changing to IL-7 supplementation 5 days post transduction. Comparable results regarding the proportion of CD90.2^+^ cells (a) and the expression levels of the CD90.2 surface marker (b), TCRβ 9c) and TCRζ within the CD90.2^+^ population were obtained, except for the steep decline in the percentage of CD90.2^+^ cells observed for cells transduced with pMY observed before. Note that without the addition of IL-7 the cells stop proliferating strongly after 5 days post-transduction and therefore the experiment was terminated at day8.

**Supplemental Figure 3:**
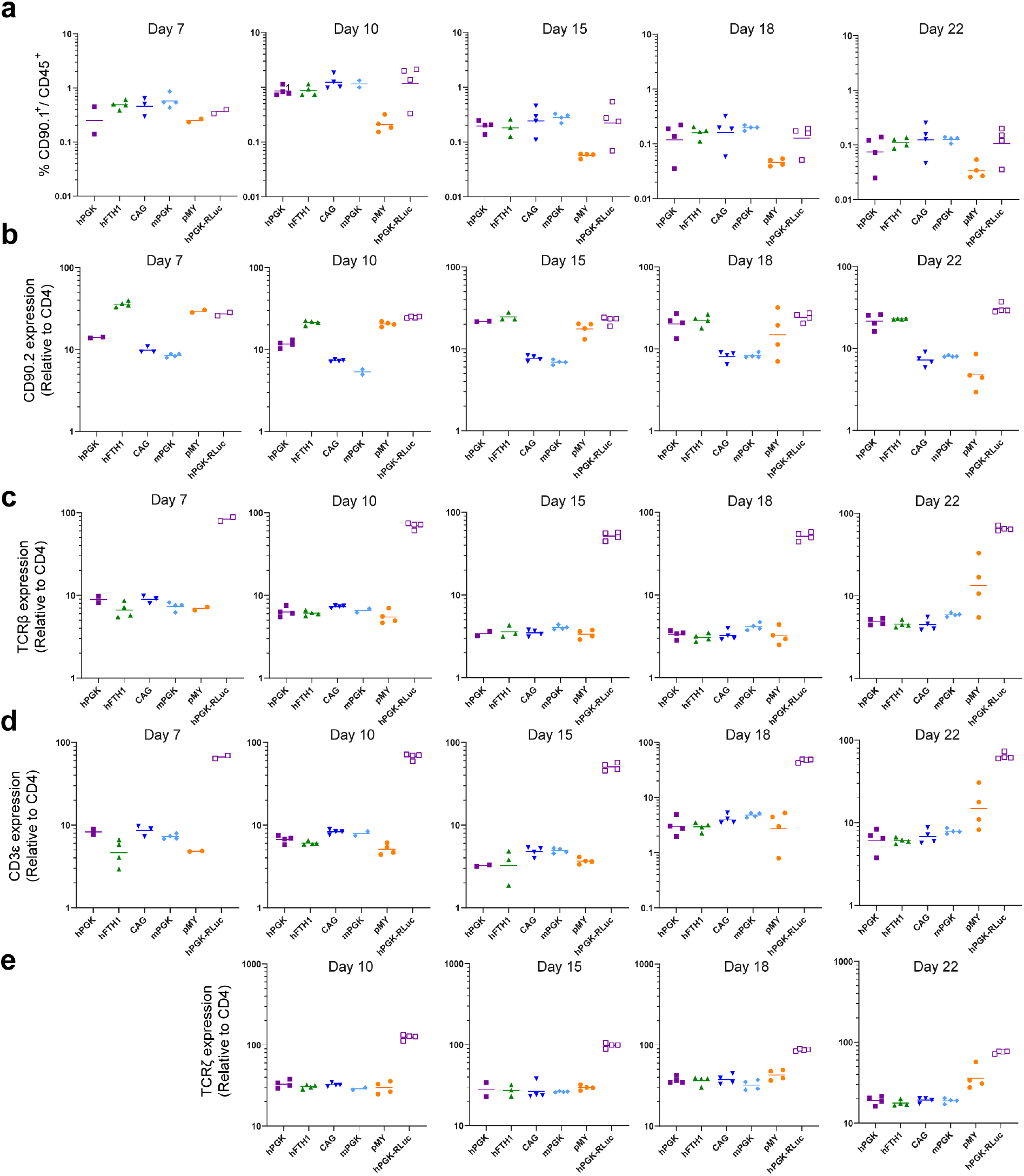
Individual values of *in vivo* vector validation in blood. Individual values corresponding to the data of figure 5a-e. Cells obtained from tail-vein bleeds at the indicated timepoints were assessed by flow cytometry. Bars depict the mean.

**Supplemental File 1: Full information on all vectors used in this study and constructed as part of the vector starter set.**

## Methods

### Molecular cloning

The pMX vector backbone was created by modifying pMXs-IRES-GFP (Cellbiolabs/Bio-connect, NL) by removing the eGFP, IRES and MCS insert by restriction with ClaI and SalI and ligating a synthetic MCS fragment containing PacI, BamHI, NotI, MluI, SphI, SbfI, HindIII and ClaI restriction sites respectively (TTAATTAACTCCGTGGATCCGGTCGTGCGGCCGCACGGAAACGCGTGGCCTGGCATGCCGCGACCCTGCAGGTTTCTGAAGCTTGAGTACATCGAT).

Similarly, the original pMY-IRES-GFP vector (Cellbiolabs/Bio-connect, NL) was modified to replace the multiple cloning site. Furthermore, the second SalI site in this vector was removed by restriction with SalI followed by 3’exonuclease digestion and blunt-end ligation, creating pMY-Empty-MCS (Addgene: #163351).

A lentiviral vector suitable for convenient subcloning between these modified pMY and pMX vectors was made by modifying lentiCRISPR v2 (Addgene: #52961) (Sanjana, Shalem, & Zhang, 2014). First, the U6 and gRNA scaffold were removed by restriction with KpnI and EcoRI followed by blunt ligation. Next, the Cas9 insert was replaced by eGFP by cloning with AgeI and BamHI. The full GFP-P2A-PuroR including the 3'WPRE and 3'LTR was shuttled to the empty pMY backbone with PacI and ApaI to allow easier modification. Here, the GFP-P2A-PuroR insert was removed by PacI and MluI and replaced with the MCS sequence with PacI and ClaI. This insert was shuttled back into the pLenti backbone with PacI and ApaI replacing the original fragment. Next the endogenous NotI and MluI sites were removed consecutively by restriction followed by 3'exonuclease digestion and blunt-end ligation, creating pLenti-EFS-MCS-WPRE (Addgene: #163362).

The different promoter sequences were obtained by synthesis (GeneArt) and were amplified by PCR before being ligated into the pMX-vector backbone with PacI and BamHI (Supplemental table 2 for primers). The murine surface marker Ly6G was codon optimized and synthesized (Geneart, supplemental methods 1). The sequence was amplified with primers Ly6Gopt_SalP2A_Fwd and Ly6Gopt_Sbf_Rev and ligated into pMY with SalI and SbfI, creating pMY-MCS-P2A-Ly6G (Addgene: #163353). Similarly, the sequences for CD90.1 and CD90.2 (also known as Thy1.1 and Thy1.2) were amplified from C7, or C57bl/6 cDNA with primers CD90.2_SalP2A_Fwd and CD90.2_Sbf_Rev and ligated into pMY, creating vectors pMY-MCS-P2A-CD90.1 (Addgene: #163354) and pMY-MCS-P2A-CD90.2 (Addgene: #163355). P2A-GFP was amplified from a previously constructed vector (pMX-CAF-Slc7a1-P2A-GFP unpublished) with primers P2A_SalI_Fwd and GFP_Sbf_Rev and was cloned into pMY with SalI and SbfI, creating vector pMY-MCS-P2A-GFP (Addgene: #163356). Similarly, puromycin and blasticidin resistance genes were amplified from existing vectors and were labeled with an HA-tag and DYK-tag respectively. To this end, the blastR cassette was amplified with primers BlastR_SalP2A_Fwd and BlastR_DYK_Sbf_Rev and the PuroR cassette with primers PuroR_SalP2A_Fwd and PuroR_HA_Sbf_Rev. These products were cloned into the pMY backbone with SalI and SbfI to create pMY-MCS-P2A-PuroHA (Addgene: #163352) and pMY-MCS-P2A-BlastDYK (Addgene: #163357) respectively.

The transmembrane and intracellular components of the CD3 complex CD3γ, CD3δ, CD3ε and TCRζ were codon optimized and synthesized on an expression construct, where the genes were separated by T2A, F2A and E2A peptides, followed by a P2A peptide and the codon optimized Ly6G surface marker (Supplemental methods 1). This construct was cut with NotI and SalI and ligated into pMY-MCS-P2A-CD90.2 to express it in frame with the CD90.2 surface marker instead of Ly6G, creating pMY-CD3-P2A-CD90.2 (Addgene: #163338). The full CD3-CD90.2 insert was further subcloned into the pMX vectors with different promoters using BamHI and HindIII (Addgene #163334-7).

The gene encoding mCherry was amplified with a GSG linker from vector pMSCV-nMCL1GFP-IRES-mCherry (Unpublished, kind gift from Chiara Montironi and Eric Eldering, Amsterdam UMC), which is a derivate of pMSCV-IRES-mCherry (Addgene #52114), with primers mCherry_Sal_GSG_Fw and mCherry_Hind_Rv. This product was cloned into pMY to create the intermediate product pMY-MCS-GSG-mCherry. This was used as a backbone to clone murine *lyz2* into, which was amplified from C57bl/6 mouse cDNA with primers Lyz2_MluI_Fwd and Lyz2_GSG_SalI_Rev, creating vector pMY-Lyz2-GSG-mCherry (Addgene: #163346). This vector was used as a backbone to replace the genes encoding the other fluorescent proteins. mTurquoise2 (Addgene: #163347), mVenus (Addgene: #163348). In parallel, GSG-eGFP was cloned in the pMY empty vector creating pMY-SalI-GSG-eGFP (Addgene: #163350). These GSG-FP fragments were amplified with primers mCherry_Sal_GSG_Fw and mCherry_Hind_Rv. The template for mTurquoise2 was pEGFP-N1-4xmts-mTurquoise2 Addgene #98819 (Chertkova et al., 2017), for mVenus it was pEGFP-C1-SYFP1 (unpublished, kind gift from Joachim Goedhart) (Goedhart et al., 2012; Kremers et al., 2006).

The MiR30 fragment was amplified from from pGIPZ-miR30-FYN (Horizon discovery) with primers miR30_Hind_Fwd and miR30_Cla_Rev and was cloned into pMY-Ly6G-P2A-PuroHA, pMY-LygG, pMY-CD90.2-Blast-DYK, or pMY-CD90.2 with HindIII and ClaI. Target sequences for antagomirs were selected with help of the genetic perturbation platform and the antagomir fragments were designed according to published guidelines for miR30 generation (Chang et al., 2013a; Dow et al., 2012). Antagomir sequences were synthesized as single stranded oligonucleotides SHC007_shRNAmiR_temp (Rluc) and Cd247A_miR_temp (CD247/TCRζ) and were amplified with primers miRE-Xho-Fwd and miRE-Eco-Rev before being inserted in the miR30 backbone with XhoI and EcoRI.

### Isolation, activation and culture of C7 cells

Transgenic C7-TCR.CD90.1 mice (Gallegos et al., 2008, 2016) were killed by administration of a sublethal dose of 0.1 ml KetMet/10 g of mouse weight (KetMed consists of 12.5 mg/ml ketamine and 30 μg/ml dexmedetomidine), followed by cervical dislocation. Spleens and lymph nodes were collected and homogenized through a 100 μm EASYstrainer cell strainer (Greiner). After washing the cells with PBS, they were resuspended in 900 μl MACS buffer and 100 μl anti-CD4 MACS bead suspension (CD4 L3T4 microbeads Miltenyi) was added and incubated on ice for 15 minutes. The suspension was centrifuged and resuspended in 1 ml MACS buffer, divided over 2 35 μm cell strainers (FALCON 5 ml round bottom tube with cell strainer cap) and spun down (500g). Cells were resuspended in 3 ml MACS buffer and were applied to a prewashed LS MACS column (Miltenyi) on a MACS magnet. The CD4^-^ fraction was collected by triple washing with 3 ml MACS buffer and collecting flow through. Afterwards, the LS column was removed from the magnet and the remaining cells were collected as CD4^+^ fraction.

Cells were counted on a CASY cell counter and the CD4^-^ cells were irradiated by exposure to a Cesium (^137^C)-source to receive 10gy. C7 cells were activated by adding 1.5*10^6^ irradiated CD4^−^ cells to 0.5*10^6^ CD4^+^ cells per ml RPMI+, supplemented with 10 ng/ml IL-12 and 5 μg/mL ESAT6_1-20_ peptide (produced by the Netherlands Cancer Institute (NKI).

CD4^+^ and non-irradiated CD4^−^ fractions were stained with flow cytometry panel 2 to assess MACS efficiency.

### Transfection and transduction

To produce ecotropic retrovirus, platinum-E (PLAT-E) cells were transiently transfected (Kitamura et al., 2003). To achieve this, PLAT-E cells were pre-cultured in IMDM medium supplemented with 10% fetal calf serum, penicillin and streptomycin. From an exponentially growing PLAT-E culture, 3*10^6^ cells were inoculated in 45 ml IMDM+ in a T225 culture flask 72 hours before transfection. On the day of transfection, PLAT-E cells were washed with 45 ml of PBS and were dissociated from the culture flask by a 5 minute incubation (37°C) with 9 ml of TrypLE reagent (Gibco), which was inactivated by adding 36 ml PBS. The PLAT-E cells were washed, resuspended in IMDM+ and filtered over a 40 μm cell strainer, after which the cells were counted with a CASY cell counter. 2.5*10^6^ PLAT-E cells were suspended in 1 ml IMDM and these cells were transfected by adding the transfection mix.

The transfection mix was made by dissolving 2 μg of the indicated vectors in combination with 0.4 μg of the helper plasmid pCL-ECO (Addgene plasmid 12371) (Naviaux et al., 1996) and for miR30 vectors 0.4 μg DGCR8 siRNA (Chang, Marran, Valentine, & Hannon, 2013b) (synthesized by Qiagen) in a total volume of 95.4 μl Opti-MEM (Gibco). After mixing of DNA by flicking tubes and a short spin, 5.6 μl P3000 reagent was added (Thermofisher). Simultaneously, 5.6 μl P3000 transfection reagent per sample was dissolved in 94.4 μl Opti-MEM, this mix was added to the DNA-containing mix and incubated for 10 minutes. After incubation, 800 μl of IMDM was added and the mix was added to the PLAT-E cells in 6-well culture dish. One day after transfection the IMDM medium was carefully removed from the transfected cells and replaced with 1.5 ml of RPMI medium containing L-glutamine 50 μM β-mercaptoethanol, 10% FBS and 10,000 U/mL penicillin/streptomycin (Referred to as RPMI+ for the rest of the methods section).

After overnight incubation, virus-containing supernatant was collected and filtered over a 0.2 μm filter. Activated C7 cells were concentrated and 4 ml of culture was resuspended in 1 ml RPMI+ containing 10 μg/mL ESAT61-20 peptide + 20 ng/mL IL-12 (Peprotech) + 20 ng/mL IL-2 (Peprotech). 1 ml of virus-containing supernatant was added to 1 ml activated C7 cells on retronectin coated 6-well culture plates and centrifuged for 2 hours on 1000x *g*. After further 3 hours of culture, 2 ml of RPMI+ containing 10 ng/ml IL-2 was added to each well. Remaining PLAT-E cells were washed with PBS and stained with flow cytometry panel 2 (below) to assess transfection efficiency.

Similar procedures were performed for the production of amphotropic retrovirus. However, PLAT-A cells (Kitamura et al., 2003) were used instead of PLAT-E cells. Since PLAT-A cells tend to dissociate easily from plates once virus production has started, culture plates were pre-treated with Poly-D-Lysine. Finally, the helper plasmid pCL-Ampho (Naviaux et al., 1996) (Novus Biologicals NBP2-29541) was used instead of pCL-Eco.

### Flow Cytometry

Plat-E cells were dissociated by TrypLE reagent (Gibco) before FACS staining while C7 cells were stained directly. All cells were first stained for viability with Fixable Viability Dye eFluor™ 780 (1:1000 in PBS) (eBioscience), before cell surface staining with the antibody combination depicted in supplemental table 3, depending on the condition. Antibody cocktails were prepared in FACS buffer (PBS containing 0.5% bovine serum albumin (BSA; Sigma) and 0.1% NaN_3_) cells were stained for 10-15 minutes at 4 °C before fixation in 2% paraformaldehyde solution for 5 minutes (Electron Microscopy Sciences). In the case of intracellular staining for TCRζ, cells were permeabilized, by incubation with Perm/Wash solution (BD Biosciences) for 5 min at 4 °C, before intracellular staining and second fixation step. Flow cytometry was performed on Canto flow cytometer (BD Bioscience) and data was analyzed using FlowJo V10 software (TreeStar).

### Confocal microscopy

C7 T cells were washed in PBS, fixed in 2% PFA for 5 minutes washed again and mounted on glass slides (ProLong Gold mounting medium without DAPI), two days and five days after transduction. Slides were imaged with a Leica SP8X confocal microscope using a 63x objective (Numerical aperture 1.4) using LasX software (Leica, version 3.5.6). Blue fluorophore mTurquoise2 was excited with a UV laser at 405 nm (Shutter 20%, Laser power 50%, Laser strength 2%) and emission was measured at 488-493 nm (HyD, Gain 100V, Offset −0.2%, Pinhole 0.7 Airy units). Other fluorophores were excited with a white-light laser (Shutter 20%, Laser power 50%, Laser strength 2%) at 488 nm for eGFP, 515 nm for mVenus and 587 nm for mCherry and emission was acquired at 500-535 nm (eGFP), 520-560 nm (mVenus), or 610-650 nm (mCherry) (HyD, Gain 600 V, Offset 0.0%, Pinhole 0.7 Airy units). Images were acquired at 10x zoom, resolution of 512×512 pixels, a line average of 4 images, bidirectional X imaging, speed setting of 600 and a distance between Z-planes of 0.2 nm. Images were deconvolved using Huygens Professional Software suite (Version 19.10) and images were 3D rendered in LasX software (Version 3.5.6).

### *In vivo* experiments

Transduced cells were split 1:2 one day after transduction by adding fresh RPMI+ containing 10 ng/ml IL-2. Two days after transduction, cells were washed and suspended in PBS to reach 1*10^7^ cells/ml. 200 μl of this solution (*i.e*. 2*10^6^ cells) was intravenously injected into the tail vein of healthy C57Bl6 mice (7 weeks old). Recipient mice were randomly distributed over 6 cages by animal caretakers and experimental groups were separated per cage after receiving C7 cells. One mouse belonging to the group transduced with CAG-containing construct did not receive the full 200 μl of C7 cells. Blood aliquots were collected in EDTA containing tubes at the indicated timepoints by tail vein bleed (Maximally 50 μl per bleed) after puncture with a 25G medical needle.

EDTA tubes containing blood samples were spun, resuspended in red blood cell lysis buffer and incubated for 2 minutes. After this lysis step, cells were resuspended in FACS buffer and kept at 4 °C until staining with flow cytometry panel 3. Please note that the first bleed at 5 days post transduction yielded insufficient cells to perform an intracellular staining and therefore no data on TCRζ-expression are available for this timepoint. Furthermore, not all samples could be analyzed at this timepoint and therefore some error bars are missing from the corresponding figure.

Recipient mice were sacrificed 22 days after T cell activation (*i.e*. 18 days after adoptive transfer) as above. 200 μl of blood was collected in EDTA tubes and analyzed as above. Lungs and spleens were harvested from all mice. Lungs were homogenized by cutting with sterile scissors in the presence of 0.5 ml digestion buffer (HBSS supplemented with 5mM CaCl_2_ and 200 U/mL collagenase IV). 2 ml digestion buffer was added to the lung fragments and transferred to 12 ml round bottom tubes, which were incubated for 30 minutes at 37 °C while shaking at 225 rotations per minute. Digested lung sample was homogenized by passing 10 times through a 19 G needle fitted to a 2 ml syringe. Homogenized tissue was filtered over a 100 μm EASYstrainer (Greiner) cell strainer, which washed with 10 ml PBS. Spleens were pooled per experimental group and were homogenized by sieving through a 100 μm EASYstrainer. Cells were resuspended in PBS and analyzed by flow cytometry as above.

## Supplemental Methods 1: Synthesized sequences

### CD3-complex optimizednotI-gamma-mfeI-T2A-delta-mluI-F2A-epsilon-SphI-E2A-zeta-SalI-P2A-Ly6G-SbfI

GCAGAACCGCGGCCGCGCCACCATGGAACAGAGAAAAGGCCTGGCCGGCCTGTTCCTGG

TTATCAGTCTGCTGCAGGGCACAGTGGCCCAGACCAACAAGGCTAAGAACCTGGTGCAG

GTGGACGGCTCTAGAGGCGACGGATCTGTGCTGCTGACATGTGGCCTGACCGACAAGACC

ATCAAGTGGCTGAAGGACGGCTCCATCATCAGCCCTCTGAACGCCACCAAGAACACCTGG

AACCTGGGCAACAACGCCAAGGACCCCAGAGGCACCTATCAGTGCCAGGGCGCCAAAGA

GACAAGCAACCCTCTGCAGGTCTACTACAGAATGTGCGAGAACTGCATCGAGCTGAACAT

CGGCACCATCAGCGGCTTCATCTTCGCCGAAGTGATCAGCATCTTCTTTCTGGCCCTGGGC

GTGTACCTGATCGCTGGACAAGATGGCGTGCGGCAGAGCAGAGCCAGCGATAAGCAGAC

ACTGCTGCAGAACGAGCAGCTGTACCAGCCTCTGAAGGACAGAGAGTACGACCAGTACA

GCCACCTCCAGGGCAACCAGCTGCGGAAGAAGGGATCTGGCCAATTGGAAGGCAGAGGC

TCTCTTCTTACATGCGGCGACGTCGAGGAAAACCCAGGACCTATGGAACACTCTGGCATC

CTGGCTAGCCTGATCCTGATTGCCGTTCTGCCTCAAGGCAGCCCCTTCAAGATCCAAGTG

ACCGAGTACGAGGACAAGGTGTTCGTGACCTGCAACACCAGCGTGATGCACCTGGATGG

CACCGTGGAAGGATGGTTCGCCAAGAACAAGACCCTGAACCTCGGCAAGGGCGTGCTGG

ACCCTAGAGGCATCTACCTGTGTAACGGCACAGAGCAGCTGGCCAAGGTGGTGTCTAGTG

TGCAGGTCCACTATCGGATGTGTCAGAACTGCGTGGAACTGGACAGCGGCACAATGGCC

GGCGTGATCTTCATCGACCTGATCGCTACCCTGCTGCTGGCACTGGGAGTGTATTGCTTCG

CTGGCCACGAGACAGGCAGACCTAGCGGAGCTGCTGAAGTTCAGGCCCTGCTGAAGAAT

GAACAGCTCTATCAGCCCCTGCGCGACAGAGAGGATACCCAGTACTCTAGACTCGGCGGC

AACTGGCCCAGAAACAAGAAATCTGGAAGCGGCACGCGTGTCAGACAGACCCTGAACTT

CGATCTGCTTAGACTGGCCGGGGACGTCGAGTCTAATCCAGGACCAATGCGGTGGAACAC

CTTCTGGGGCATCCTGTGTCTGTCTCTGCTGGCTGTGGGCACCTGTCAGGATGACGCTGAG

AACATCGAGTATAAGGTGTCCATCTCCGGCACCAGCGTCGAGCTGACTTGTCCTCTGGAC

TCCGACGAGAACCTGAAGTGGGAGAAGAACGGCCAAGAGCTGCCTCAGAAGCACGACAA

GCACCTGGTGCTGCAGGACTTCAGCGAGGTGGAAGATAGCGGCTACTACGTGTGCTACAC

CCCTGCCAGCAACAAGAACACATACCTGTACCTGAAGGCTCGCGTGTGCGAGTACTGTGT

CGAGGTGGACCTGACAGCCGTGGCTATCATCATCATCGTGGACATCTGCATCACCCTGGG

CCTGCTGATGGTCATCTACTACTGGTCCAAGAACCGGAAGGCCAAGGCCAAGCCTGTGAC

AAGAGGAACCGGCGCTGGAAGCAGACCAAGAGGCCAGAACAAAGAAAGACCTCCTCCT

GTGCCTAATCCTGACTACGAGCCCATCCGGAAGGGCCAGAGAGATCTGTACTCTGGCCTG

AACCAGAGGGCCGTGGGTTCTGGCGCATGCCAGTGTACCAACTATGCTCTCCTGAGACTC

GCAGGCGACGTTGAGAGTAATCCAGGGCCTATGAAGTGGAAAGTGTCTGTGCTGGCCTGC

ATCCTGCATGTTCGATTCCCTGGCGCTGAGGCCCAGTCTTTTGGACTGCTGGACCCCAAGC

TGTGCTACCTGCTGGACGGCATTCTGTTTATTTATGGCGTGATCATCACCGCTCTGTACCT

GCGGGCCAAGTTCAGCAGAAGCGCTGAGACAGCTGCCAATCTGCAGGACCCTAACCAGC

TGTACAACGAGCTGAATCTGGGGCGCAGAGAAGAGTACGATGTGCTGGAAAAGAAGAGA

GCCAGAGATCCCGAGATGGGCGGCAAACAGCAGAGAAGGCGGAATCCTCAAGAAGGCG

TGTACAACGCCCTGCAGAAAGATAAGATGGCCGAGGCCTACAGCGAGATCGGCACAAAG

GGCGAACGCAGAAGAGGCAAGGGACACGATGGACTGTACCAGGGCCTGTCCACAGCCAC

AAAGGACACATACGATGCCCTGCACATGCAGACACTGGCCCCTAGAGGCAGCGGCGTCG

ACGCCACAAACTTCAGCCTGCTGAGACAGGCTGGCGACGTGGAAGAGAATCCTGGACCT

ATGGACACCTGTCATATCGCCAAGAGCTGCGTGCTGATCCTGCTGGTGGTTCTGCTGTGTG

CCGAGCGAGCACAGGGACTGGAGTGCTACAACTGTATCGGCGTGCCACCTGAGACAAGC

TGCAACACCACCACCTGTCCTTTCAGCGACGGCTTCTGTGTGGCCCTGGAAATCGAAGTG

ATCGTGGACAGCCACCGCAGCAAAGTGAAGTCCAACCTGTGCCTGCCTATCTGCCCCACC

ACACTGGACAACACCGAGATCACAGGCAACGCCGTGAACGTGAAAACCTACTGCTGCAA

AGAGGACCTCTGCAACGCCGCTGTTCCAACAGGCGGAAGCTCTTGGACAATGGCTGGCGT

GCTGCTGTTCAGCCTGGTGTCTGTTCTGCTGCAGACCTTCCTGTGACCTGCAGGGGATGCA

T

### hPGK

TTAATTAACGGGGTTGGGGTTGCGCCTTTTCCAAGGCAGCCCTGGGTTTGCGCAGGGACG

CGGCTGCTCTGGGCGTGGTTCCGGGAAACGCAGCGGCGCCGACCCTGGGTCTCGCACATT

CTTCACGTCCGTTCGCAGCGTCACCCGGATCTTCGCCGCTACCCTTGTGGGCCCCCCGGCG

ACGCTTCCTGCTCCGCCCCTAAGTCGGGAAGGTTCCTTGCGGTTCGCGGCGTGCCGGACG

TGACAAACGGAAGCCGCACGTCTCACTAGTACCCTCGCAGACGGACAGCGCCAGGGAGC

AATGGCAGCGCGCCGACCGCGATGGGCTGTGGCCAATAGCGGCTGCTCAGCAGGGCGCG

CCGAGAGCAGCGGCCGGGAAGGGGCGGTGCGGGAGGCGGGGTGTGGGGCGGTAGTGTG

GGCCCTGTTCCTGCCCGCGCGGTGTTCCGCATTCTGCAAGCCTCCGGAGCGCACGTCGGC

AGTCGGCTCCCTCGTTGACCGAATCACCGACCTCTCTCCCCAGGGATCC

### mPGK

GCAGAACCTTAATTAAAAATTCTACCGGGTAGGGGAGGCGCTTTTCCCAAGGCAGTCTGG

AGCATGCGCTTTAGCAGCCCCGCTGGGCACTTGGCGCTACACAAGTGGCCTCTGGCCTCG

CACACATTCCACATCCACCGGTAGGCGCCAACCGGCTCCGTTCTTTGGTGGCCCCTTCGCG

CCACCTTCTACTCCTCCCCTAGTCAGGAAGTTCCCCCCCGCCCCGCAGCTCGCGTCGTGCA

GGACGTGACAAATGGAAGTAGCACGTCTCACTAGTCTCGTGCAGATGGACAGCACCGCT

GAGCAATGGAAGCGGGTAGGCCTTTGGGGCAGCGGCCAATAGCAGCTTTGCTCCTTCGCT

TTCTGGGCTCAGAGGCTGGGAAGGGGTGGGTCCGGGGGCGGGCTCAGGGGCGGGCTCAG

GGGCGGGGCGGGCGCCCGAAGGTCCTCCGGAGGCCCGGCATTCTGCACGCTTCAAAAGC

GCACGTCTGCCGCGCTGTTCTCCTCTTCCTCATCTCCGGGCCTTTCGACGGATCCGGATGC

AT

### hFTH1

TTAATTAATCCGCCAGAGCGCGCGAGGGCCTCCACCGGCCGCCCCTCCCCCACAGCAGGG

GCGGGGTCCCGCGCCCACCGGAAGGAGCGGGCTCGGGGCGGGCGGCGCTGATTGGCCGG

GGCGGGCCTGACGCCGACGCGGCTATAAGAGACCACAAGCGACCCGCAGGGCCAGACGT

TCTTCGCCGAGAGTCGTCGGGGTTTCCTGCTTCAACAGTGCTTGGACGGAACCCGGCGCT

CGTTCCCCACCCCGGCCGGCCGCCCATAGCCAGCCCTCCGTCACCTCTTCACCGCACCCTC

GGACTGCCCCAAGGCCCCCGCCGCCGCTCCAGCGCCGCGCAGCCACCGCCGCCGCCGCCG

CCTCTCCTTAGTCGCCGCCGGATCC

### CAG

TTAATTAATCGACATTGATTATTGACTAGTTATTAATAGTAATCAATTACGGGGTCATTAG

TTCATAGCCCATATATGGAGTTCCGCGTTACATAACTTACGGTAAATGGCCCGCCTGGCT

GACCGCCCAACGACCCCCGCCCATTGACGTCAATAATGACGTATGTTCCCATAGTAACGC

CAATAGGGACTTTCCATTGACGTCAATGGGTGGACTATTTACGGTAAACTGCCCACTTGG

CAGTACATCAAGTGTATCATATGCCAAGTACGCCCCCTATTGACGTCAATGACGGTAAAT

GGCCCGCCTGGCATTATGCCCAGTACATGACCTTATGGGACTTTCCTACTTGGCAGTACAT

CTACGTATTAGTCATCGCTATTACCATGGGTCGAGGTGAGCCCCACGTTCTGCTTCACTCT

CCCCATCTCCCCCCCCTCCCCACCCCCAATTTTGTATTTATTTATTTTTTAATTATTTTGTGC

AGCGATGGGGGCGGGGGGGGGGGGGGCGCGCGCCAGGCGGGGCGGGGCGGGGCGAGGG

GCGGGGCGGGGCGAGGCGGAGAGGTGCGGCGGCAGCCAATCAGAGCGGCGCGCTCCGA

AAGTTTCCTTTTATGGCGAGGCGGCGGCGGCGGCGGCCCTATAAAAAGCGAAGCGCGCG

GCGGGCGGGAGTCGCTGCGTTGCCTTCGCCCCGTGCCCCGCTCCGCGCCGCCTCGCGCCG

CCCGCCCCGGCTCTGACTGACCGCGTTACTCCCACAGGTGAGCGGGCGGGACGGCCCTTC

TCCTCCGGGCTGTAATTAGCGCTTGGTTTAATGACGGCTCGTTTCTTTTCTGTGGCTGCGT

GAAAGCCTTAAAGGGCTCCGGGAGGGGGATCC

### Ly6Gopt

GCAGAACCCCTGCAGGCCACCATGGACACCTGTCATATCGCCAAGAGCTGCGTGCTGATC

CTGCTGGTGGTTCTGCTGTGTGCCGAGCGAGCACAGGGACTGGAGTGCTACAACTGTATC

GGCGTGCCACCTGAGACAAGCTGCAACACCACCACCTGTCCTTTCAGCGACGGCTTCTGT

GTGGCCCTGGAAATCGAAGTGATCGTGGACAGCCACCGCAGCAAAGTGAAGTCCAACCT

GTGCCTGCCTATCTGCCCCACCACACTGGACAACACCGAGATCACAGGCAACGCCGTGAA

CGTGAAAACCTACTGCTGCAAAGAGGACCTCTGCAACGCCGCTGTTCCAACAGGCGGAA

GCTCTTGGACAATGGCTGGCGTGCTGCTGTTCAGCCTGGTGTCTGTTCTGCTGCAGACCTT

CCTGTGAGCATGCCAATTCTCATCGATTGCATTGGGTCGACGGATGCAT

---

**Supplemental table 1:**
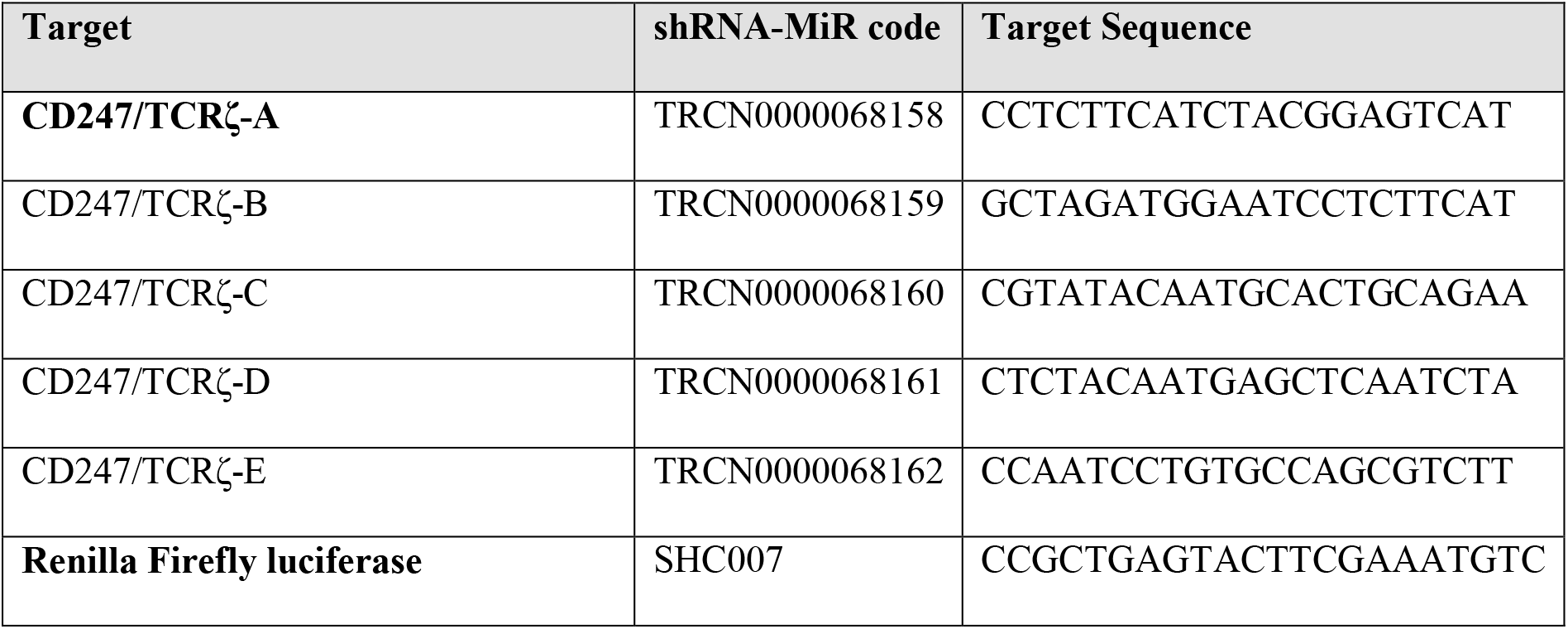
shRNA-miRs used in this study. The antagomir target sequences were derived from the genetic perturbation platform of the Broad Institute (https://portals.broadinstitute.org/gpp/public/). Bold sequences were selected and used throughout the manuscript.

**Supplemental table 2:**
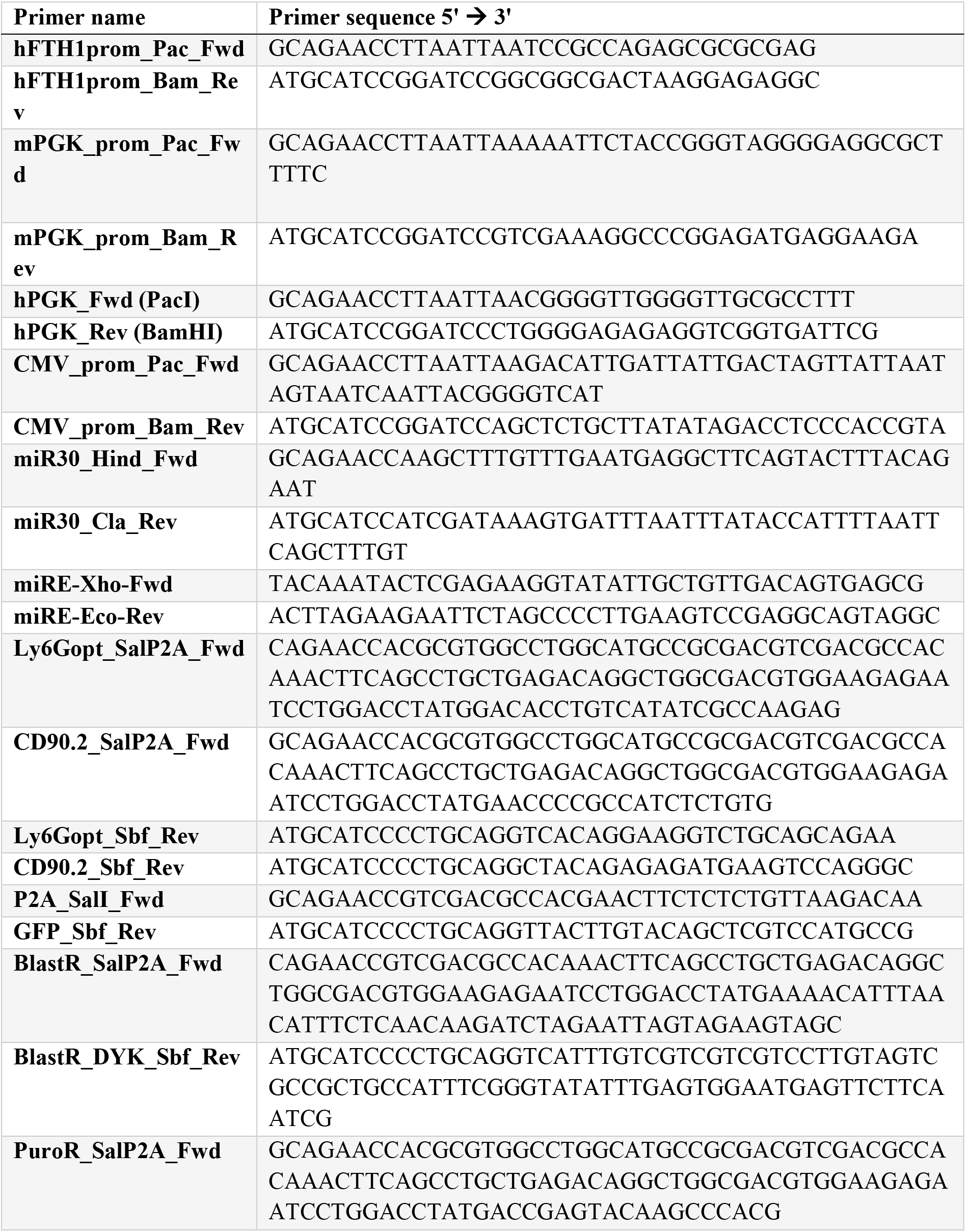
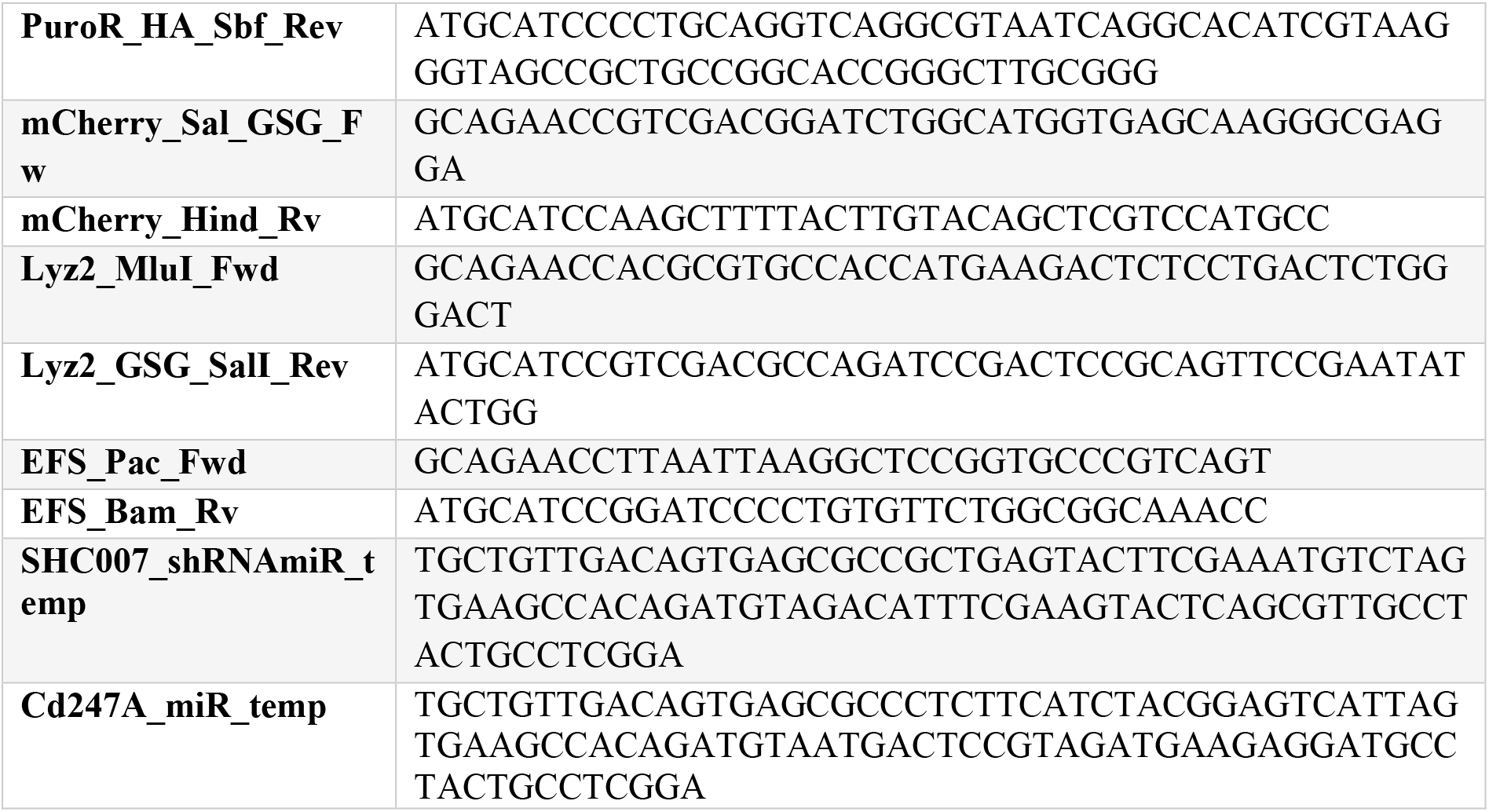
Primers used in this study.

**Supplemental Table 3:** Antibody combinations used for flow cytometry staining. Details on the individual antibodies can be found in Supplemental table 4. *: Intracellular stain performed after fixation and permeabilization. ^#^: Marker specific staining was used where opportune.

|  | Transfection<br>(Panel 1) | MACS<br>(Panel 2) | *In*<br>*vitro/vivo*<br>(Panel 3) |
| --- | --- | --- | --- |
| FITC | Ly6G<sup>#</sup> | B220 | CD3ε |
| PE |  | CD62L | TCRζ* |
| PE/Dazzle594 |  | CD45 | CD45 |
| PerCP/eFluor710 |  | CD11a | CD90.2 |
| PE/Cy7 |  | CD4 | CD4 |
| APC | CD90.2<sup>#</sup> | TCRβ | TCRβ |
| AlexaFluor700 |  | CD90.1 | CD90.1 |
| eFluor 780 | Viability | Viability | Viability |

**Supplemental Table 4:**
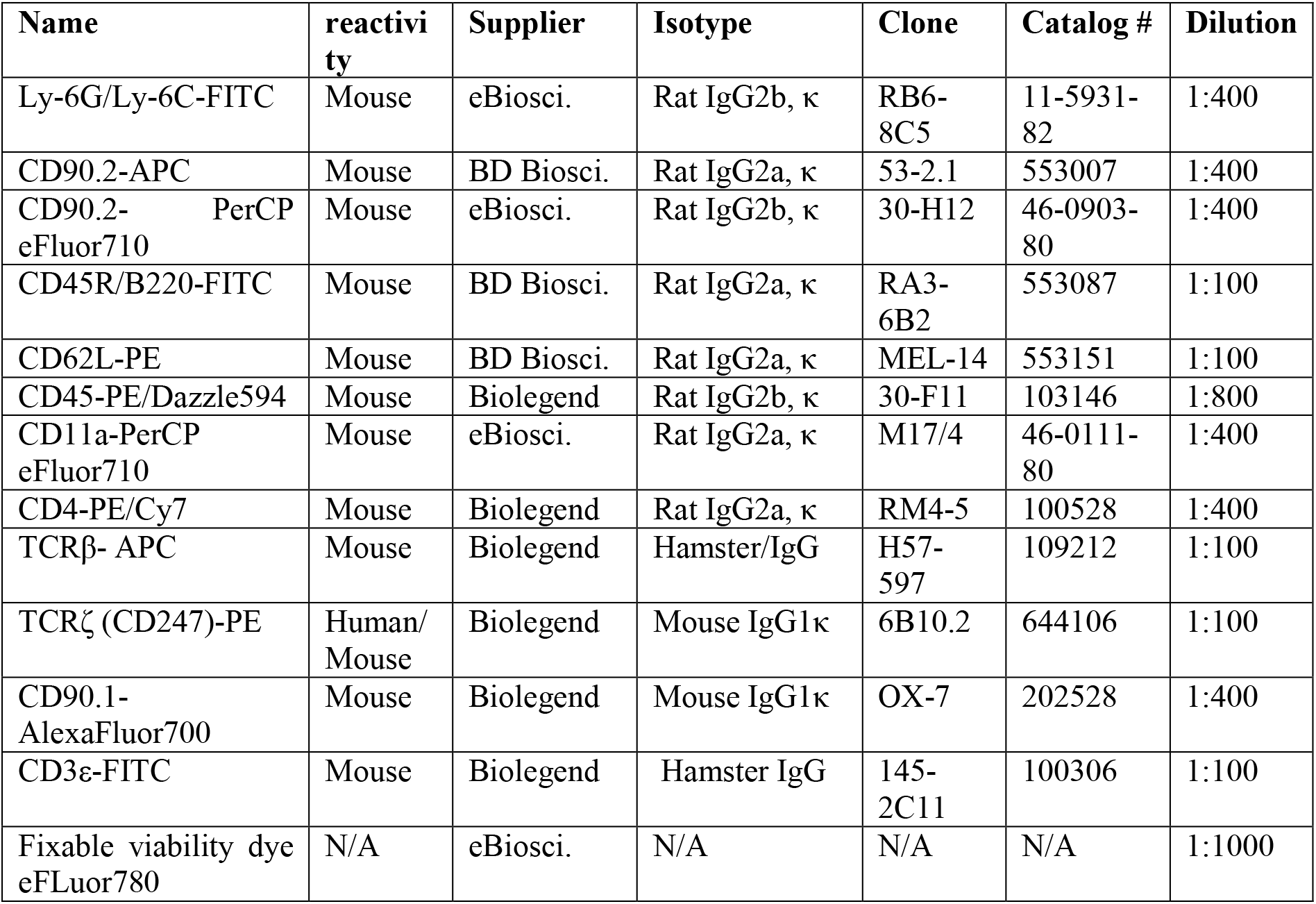
Antibody clones, suppliers and dilutions used in this study.

| Name | reactivity | Supplier | Isotype | Clone | Catalog # | Dilution |
| --- | --- | --- | --- | --- | --- | --- |
| Ly-6G/Ly-6C-FITC | Mouse | eBiosci. | Rat IgG2b, κ | RB6-8C5 | 11-5931-82 | 1:400 |
| CD90.2-APC | Mouse | BD Biosci. | Rat IgG2a, κ | 53-2.1 | 553007 | 1:400 |
| CD90.2-PerCP eFluor710 | Mouse | eBiosci. | Rat IgG2b, κ | 30-H12 | 46-0903-80 | 1:400 |
| CD45R/B220-FITC | Mouse | BD Biosci. | Rat IgG2a, κ | RA3-6B2 | 553087 | 1:100 |
| CD62L-PE | Mouse | BD Biosci. | Rat IgG2a, κ | MEL-14 | 553151 | 1:100 |
| CD45-PE/Dazzle594 | Mouse | Biolegend | Rat IgG2b, κ | 30-F11 | 103146 | 1:800 |
| CD11a-PerCP eFluor710 | Mouse | eBiosci. | Rat IgG2a, κ | M17/4 | 46-0111-80 | 1:400 |
| CD4-PE/Cy7 | Mouse | Biolegend | Rat IgG2a, κ | RM4-5 | 100528 | 1:400 |
| TCRβ- APC | Mouse | Biolegend | Hamster/IgG | H57-597 | 109212 | 1:100 |
| TCRζ (CD247)-PE | Human/<br>Mouse | Biolegend | Mouse IgG1κ | 6B10.2 | 644106 | 1:100 |
| CD90.1-AlexaFluor700 | Mouse | Biolegend | Mouse IgG1κ | OX-7 | 202528 | 1:400 |
| CD3ε-FITC | Mouse | Biolegend | Hamster IgG | 145-2C11 | 100306 | 1:100 |
| Fixable viability dye eFluor780 | N/A | eBiosci. | N/A | N/A | N/A | 1:1000 |

